# Functional Morphology of Endurance Swimming Performance and Gait Transition Strategies in Balistoid Fishes

**DOI:** 10.1101/446526

**Authors:** Andrew B. George, Mark W. Westneat

## Abstract

Triggerfishes and filefishes (Balistoidea) use balistiform locomotion to power slow steady swimming with their dorsal and anal fins and transition to a gait dominated by body and caudal fin (BCF) kinematics at high speeds. Fin and body shapes are predicted to be strong determinants of swimming performance and the biomechanics of gait transitions. The goal of this study was to combine morphometrics and critical swimming tests to explore relationships between balistoid fin and body shapes and swimming performance in a phylogenetic context in order to understand the evolution and diversification of the balistiform swimming mode. Among the 13 species of balistoid fishes examined, fishes with high aspect ratio fins tended to achieve higher critical swimming speeds than fishes with low aspect ratio fins. Species with long, large median fins and wide caudal peduncles tended to use the balistiform gait alone for a larger percentage of their total critical swimming speed than fishes with short, small median fins and narrow caudal peduncles. Fishes on both ends of the aspect ratio spectrum achieved higher swimming speeds using the balistiform gait alone than fishes with median fins of intermediate aspect ratios. Each species is specialized for taking advantage of one gait, with balistiform specialists possessing long, large median fins capable of the large power requirements of swimming at high speeds using the median fins alone, while BCF specialists possess short, small median fins, ill-suited for powering high-speed balistiform locomotion, but narrow caudal peduncles capable of efficient caudal fin oscillations to power high-speed locomotion.

**Summary Statement:** Geometric morphometrics reveal that fin and body shapes are good predictors of endurance swimming performance and gait transition strategies of triggerfishes and filefishes.

## INTRODUCTION

Fishes employ a wide variety of biomechanically distinct swimming modes to power aquatic locomotion, and this functional diversity is often reflected by the morphology of the fishes. Accordingly, fish swimming modes are defined based on the parts of the body involved in forward thrust production during steady swimming (Breder, 1926; Webb, 1984; reviewed in Sfakiotakis et al., 1999). The two major categories of classically defined swimming modes are body/caudal fin (BCF) locomotion and median/paired fin (MPF) locomotion. BCF swimmers contract their axial musculature in order to undulate sinusoidal body bending waves along the body, or oscillate their caudal fins. Conversely, MPF swimmers rely on undulations or oscillations of their median or paired fins for propulsion, while holding their bodies and caudal fins steady at most speeds. The reliance of fishes on particular anatomical features for locomotion has led to extensive research aimed at understanding important trends between fin shape, body shape and swimming performance (Nursall, 1958; Walker and Westneat, 2002; Wainwright et al., 2002; Rouleau et al. 2010; Xin and Wu, 2013). These studies (Nursall, 1958; Walker and Westneat, 2002; Wainwright et al., 2002) have revealed a widespread correlation between increasing aspect ratio (AR) of the fins involved in propulsion and increasing swimming performance across a variety of swimming modes. Aspect ratio is essentially a measure of how “wing-like” an airfoil is, and in the context of fish fins it is typically defined as the span of the fin squared, divided by the surface area of the fin (Nursall 1958; Lighthill, 1970). The theory behind high AR fins leading to high endurance swimming performance is based in hydrodynamic efficiency. Specifically, high AR fins reduce the production of destabilizing tip vortices and experience decreased drag due to lift along edge of the fin (Bushnell and Moore, 1964; Vogel, 1994). However, relationships between fin ARs and fish swimming hydrodynamics and performance have generally been examined theoretically (Lighthill, 1970; Karpouzian et al., 1990; Xin and Wu, 2013) or experimentally (Drucker and Jensen, 1996; Walker and Westneat, 2002; Wainwright et al., 2002; Fulton and Bellwood, 2004) in fishes that power locomotion with oscillatory fin kinematics, leaving these relationships largely unexplored in the context of undulatory fins.

An important characteristic of virtually all MPF swimmers, and a central focus of this study, is the fact that they undergo a gait transition with increasing speed from their respective steady MPF gait to an unsteady burst-and-glide BCF gait (Whoriskey and Wooton, 1987; Wright, 2000; Korsmeyer et al., 2002; Walker and Westneat, 2002; Cannas et al., 2006; Hale et al., 2006; Svendsen et al. 2010; Feilich, 2017). The gait transition from MPF to BCF propulsion requires the recruitment of axial body musculature and caudal fin oscillations, suggesting that body and caudal fin shape might play an important role in determining swimming efficiency and performance of MPF swimmers. Despite extensive research on pectoral fin shape and gait transitions in fishes (Drucker and Jensen, 1996; Walker and Westneat, 2002), the relationships between body and caudal fin morphometrics and endurance swimming performance of MPF swimmers have not been previously explored. Thus, a central goal of this study was to explore patterns of fish lateral profile morphometrics, including the dorsal, anal and caudal fins as well as body shape, and their associations with swimming performance and gait transitions.

Fishes in the superfamily Balistoidea are an ideal system in which to test hypotheses about morphology and swimming performance across a range of MPF kinematic patterns and gait transition strategies. This monophyletic superfamily is made up of 42 triggerfish species (Balistidae) and 107 filefish species (Monacanthidae), all of which power slow forward locomotion using their dorsal and anal fins while holding their bodies and caudal fins steady in the balistiform swimming mode. Despite this shared swimming mode, balistoid fishes possess a wide range of morphologies from the deep-bodied *Balistes* and *Brachaluteres* genera to the elongate *Oxymonacanthus* and *Anacanthus* genera (Dornburg et al., 2011; Hutchins and Swainston, 1985; Froese and Pauly, 2018). Additionally, balistoid fishes possess median fins spanning a morphological continuum from high AR, posteriorly-tapering fins of *Odonus niger* and *Canthidermis sufflamen* to low AR, rectangular fins of *Oxymonacanthus* and *Aluterus* species (Wright, 2000; Dornburg et al., 2011; Froese and Pauly, 2018). Coupled with this morphological diversity, balistoid fishes also lie on a kinematic continuum from swimming powered by highly oscillatory, flapping median fin kinematics to highly undulatory, wave-like median fin kinematics (Wright, 2000; Lighthill and Blake 1990). Furthermore, balistoid fishes with high AR median fins tend to use more oscillatory fin kinematics, while fishes with low AR median fins utilize more undulatory fin kinematics (Wright, 2000). Balistoid fishes undergo a gradual gait transition with increasing speed from balistiform locomotion alone at low speeds, to a gait that includes both balistiform locomotion *and* a small BCF contribution at intermediate speeds, and finally to a gait dominated by burst-and-glide BCF locomotion at their fastest speeds (Wright, 2000, Korsmeyer et al., 2002). Wright (2000) examined relationships between morphology and swimming performance of balistoid fishes, and discovered that triggerfishes with higher AR median fins were capable of increased endurance swimming performance and higher gait transition speeds compared to triggerfishes with lower AR fins. To build on this prior work, a second major goal of the present study was to increase the sampling of triggerfishes, add a set of filefishes, and interpret morphometrics and swimming performance datasets within a well-resolved phylogeny of the Balistoidea.

Recent advances in our understanding of balistoid phylogenetics (Dornburg et al., 2008, 2011; Santini et al., 2013; McCord and Westneat, 2016) and the development of rigorous phylogenetic comparative methods have allowed us to account for phylogenetic structure in analyses of balistoid functional morphology relationships. Additionally, the advancement of geometric morphometric techniques now allows for fine-scale analyses of morphological diversity (Dornburg et al., 2011; Feilich, 2016) beyond linear measurements and ratio calculations (Wright, 2000). Using these methods, we set out to quantify important axes of shape variation within balistoid fishes in a phylogenetic context. The primary goals of this study were thus to use endurance swimming performance tests, geometric morphometrics and phylogenetic comparative methods to test functional hypotheses between balistoid fin and body shapes and swimming performance in order to better understand the evolution and subsequent functional diversification of the unique balistiform swimming mode.

## MATERIALS AND METHODS

### Species selection and care

Swimming performance data were analyzed for eight triggerfish and five filefish species (Fig. 1). We combined swimming performance data for 7 triggerfish species (*Balistapus undulatus* Park 1797, *Balistoides conspicillum* Bloch and Schneider 1801, *Melichthys vidua* Richardson 1845, *Odonus niger* Rüppell 1836, *Rhinecanthus aculeatus* Linnaeus 1758, *Sufflamen chrysopterum* Bloch and Schneider 1801, and *Xanthichthys auromarginatus* Bennett 1832) and one filefish species (*Cantherhines macrocerus* Hollard 1853) from Wright (2000), with new performance measures from one additional triggerfish species (*Sufflamen bursa* Bloch and Schneider 1801) and four additional filefish species (*Acreichthys tomentosus* Linnaeus 1758, *Oxymonacanthus longirostris* Bloch and Schneider 1801, *Paraluteres prionurus* Bleeker 1851 and *Pervagor janthinosoma* Bleeker 1854). Data from *Pseudobalistes fuscus* Bloch and Schneider 1801 (Wright, 2000) were not included in our analyses, as swimming data were only available for two individuals. Swimming data from a total of 54 individuals were included in this swimming study with an average of 4.2 individuals per species (range: 3 – 5). All individuals were post-juveniles with standard lengths ranging from 4.8 to 10.9 cm (avg. = 7.76 cm).

**Fig. 1:**
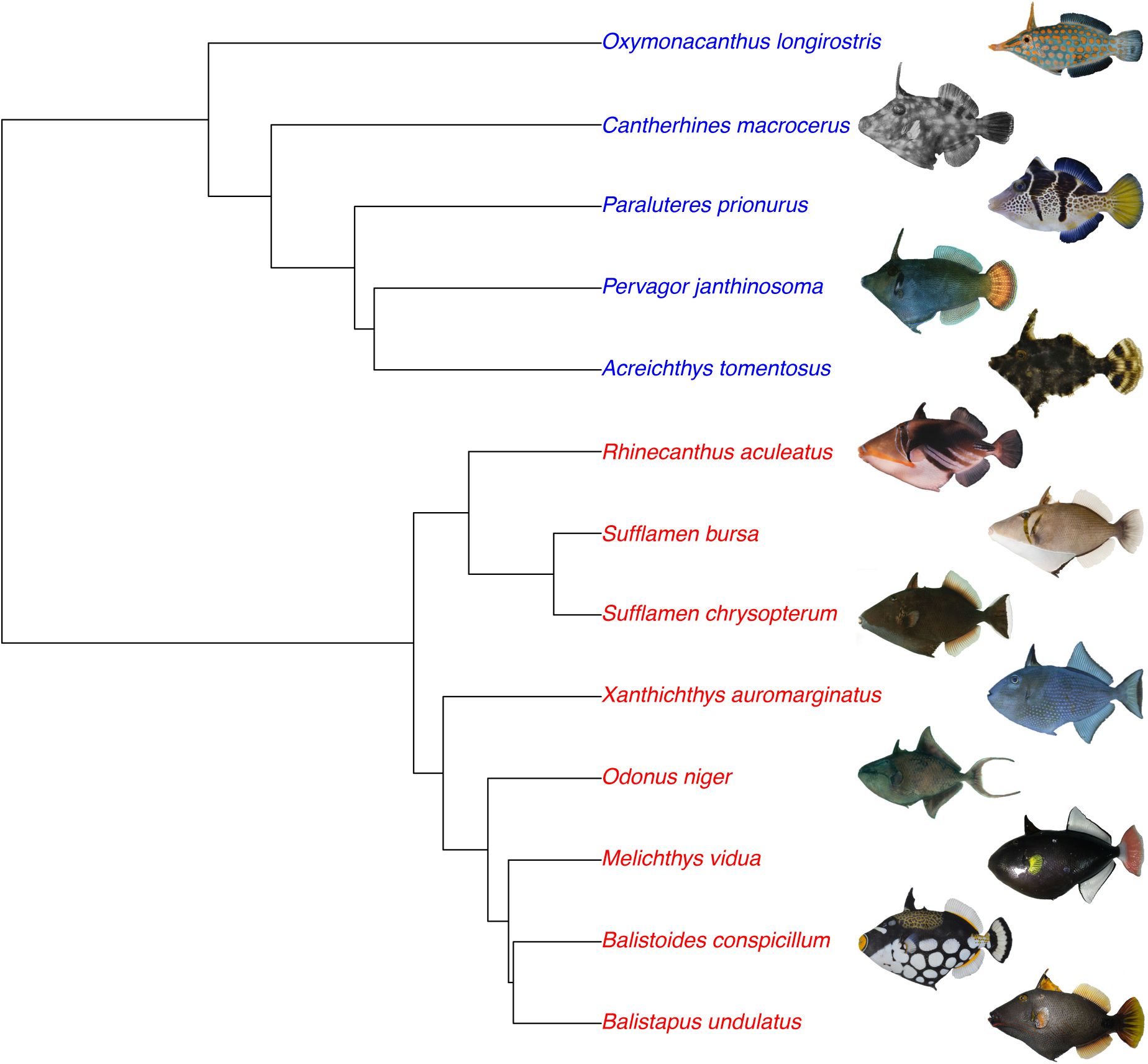
Phylogeny of the balistoid species used in this study. Species are color-coded with triggerfishes (Balistidae) in red and filefishes (Monacanthidae) in blue. Photo credits: *A. tom., B.con., P. pri*. (KPM-NR 53324, KPM-NR 45205, KPM-NR 57283; photos by Hiroshi Senou); *B. und., M. vid*. (George and Westneat); *C. mac., O. lon., P. jan., R. acu., S. chr*. (John E. Randall), *O. nig*. (Rick Winterbottom), *X. aur*. (ROM 40935, photo by Rick Winterbottom), *S. bur*. (USNM 439728; photo by Jeffrey T. Williams, Copyright 2006 Moorea Biocode, Smithsonian Institute). Phylogeny trimmed from McCord and Westneat, 2016.

Prior to swimming performance tests, all fishes were housed in separate tanks connected through a 1200-liter saltwater flow-through system. The artificial seawater in this system was maintained at a temperature of 24 ± 1°C and specific gravity of 1.024 ± 0.001. All fishes were fed freeze dried krill, fish flakes and pellets with the exception of corallivorous *O. longirostris*, which were provided live brown *Acropora spp*. coral. All fishes were deprived of food for 24 hours before swimming tests in order to control for metabolism (Alsop and Wood, 1997). All animal care protocols were approved by University of Chicago IACUC 72365.

### Endurance swimming performance tests

Fish swimming performance is typically measured using a flow tank, in which fishes are forced to swim against a controlled flow. The standard metric for assessing a fish’s endurance swimming performance is critical swimming speed (Ucrit) defied as

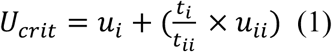

where *u_i_* is the penultimate velocity (in cm s^-1^) reached by the fish, *t_i_* is the time (in minutes) that the fish swam at the highest reached velocity before exhaustion, *t_ii_* is the length (in minutes) of each velocity increment, and *u_ii_* is the prescribed velocity step increment (in cm s^-1^) (Brett, 1964). The results of critical swimming performance tests in fishes are highly dependent on the chosen magnitude and timing of the velocity step increment (*u_i_* and *t_i_*, respectively), complicating direct comparisons of critical swimming performance between studies (Farlinger and Beamish, 1977; reviewed in Kolok, 1999). In order to ensure reliable comparisons between balistoid U_crit_ data gathered from the literature (Wright, 2000) and those measured in this study, we followed the critical swimming protocol of Wright (2000) exactly. Specifically, the length of each fish was quickly measured and recorded during the transfer of the fishes from their holding tanks to the flow tank. This measurement was used to calculate the length-specific velocity increments (*u_ii_*) used in the swimming tests. All fishes then performed a critical swimming test consisting of a two-hour acclimation period in which fishes swam in the flow tank at a low velocity of 0.5 - 1 total lengths (TL) s^-1^, followed by a stepwise increase in flow velocity of approximately 0.5 TL s^-1^ (*u_ii_*) (fork lengths for *O. niger*) every 15 minutes (*t_ii_*) until the fishes were exhausted, as evidenced by their inability to remove themselves from the downstream grate for greater than 30 seconds (Wright, 2000). All swimming tests were conducted in the same custom-made flow tank used by Wright (2000), with a working section with dimensions 25 cm × 33 cm × 104 cm. The working section was subdivided length-wise into three equal partitions (25 cm × 33 cm × 32 cm) using plastic “egg-crate” barriers as collimators between partitions. A thin sheet of acrylic was bent into a half-pipe shape and inserted into each subdivision of the working section in order to keep the fishes from avoiding swimming by wedging themselves in corners using their erectable dorsal spines and ventral keels. In a few cases the critical swimming performance of two or three individuals was determined at the same time by restricting each individual to its own separate partition of the flow tank. In such cases, exhausted fishes were quickly removed from their partition using a dipnet, and the remaining fishes continued the swimming test uninterrupted. Wright (2000) calibrated the flow tank velocity before the swimming trials by filming and digitizing the downstream motion of suspended particles over a range of speeds. In this study, flow speed was measured and adjusted during each swimming trial in real-time using a Höntzsch Instruments flow sensing probe (HFA serial no: 843; Waiblingen, Germany).

Critical swimming speed provides a measure of the total sustained swimming performance limits regardless of swimming gait used. In order to investigate the endurance swimming limits of balistiform locomotion alone (swimming powered by median fins only), gait transition data were recorded for each fish during the critical swimming trials. Due to the gradual nature of gait transitions from balistiform locomotion to BCF locomotion, Wright (2000) defined two gait transition speeds: U_t low_ and U_t high_. U_t low_ was defined as the speed interval during which the first signs of gait transition are evident, as indicated by occasional use of the caudal fin. This gait is characterized by steady balistiform locomotion *plus* occasional short BCF-powered bursts (*balistiform + BCF*). As U_t low_ is the speed at which fishes no longer power locomotion using the median fins alone, this speed can be considered the upper limit of swimming speed accomplished using balistiform locomotion alone. U_t high_ is defined as the speed interval during which the fish is no longer able to maintain a steady position in the flow tank using the median fins alone, as evidenced by use of the caudal fin every 10 seconds or less. This gait is characterized by frequent, large BCF-powered bursts followed by unsteady median fin powered locomotion as the fish glides downstream and prepares for the next BCF burst. Using this definition, it is possible for U_t high_ to be greater than U_crit_ if this gait transition occurs during the same velocity increment in which the fish becomes exhausted.

Following swimming performance tests, U_crit_, U_t low_, and U_t high_ were calculated for each individual. In order to control for the effect of fish size on swimming performance, these swimming performance metrics were expressed in terms of standard lengths per second (SL s^-1^) rather than raw speed (cm s^-1^) for all subsequent analyses. Finally, we calculated the percentage of the total critical swimming speed in which each fish swam using its median dorsal and anal fins only (*percent balistiform locomotion*) as…

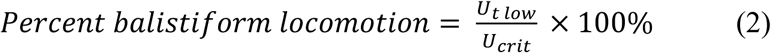

### Quantifying morphology

Following swimming experiments, fishes were euthanized with tricaine sulfonate salt (MS-222) and photographed for morphometric analyses. All fishes included in Wright’s (2000) swimming performance study were formalin-fixed and stored in 70% ethanol at the Field Museum of Natural History (FMNH) in 2000. Due to the fixation process, the fins of many of these specimens had become rigid in an unnatural position, but were otherwise in good condition. In order to spread the fins out to their natural positions for morphometric analyses, we soaked the preserved fishes in a trypsin solution for 72 hours. These fishes, along with the individuals tested in our swimming trials, were laid flat on their sides, pinned out with fins fully extended and photographed with a ruler for morphometric analyses. In order to increase the sample size for morphometric analyses, photographs of museum and aquarium trade specimens within 25% of the length range of the individuals used in the swimming experiments were included, resulting in a total of 102 individuals included in morphometric analyses (See table S1 for sample sizes of individual morphological datasets).

In order to quantify morphological diversity of fin and body shapes, a total of 109 digital landmarks were placed along the fins and bodies of the fishes using the R package *StereoMorph* (Olsen and Westneat, 2015) (Fig. 2). Twenty-six landmarks were manually placed along the fins and bodies of each fish using the *landmarks* function in *StereoMorph* (white circles and blue triangles in Fig. 2), and the dorsal, anal, and caudal fins were outlined using the *curves* function in StereoMorph (blue lines in Fig. 2). The *curves* function interprets and returns the position of landmarks along a digitized Bezier curve. Curves were digitized along the anterior, dorsal, posterior, and ventral surface of each fin, resulting in four independent curves along each fin. Following digitization, landmarks placed along the fins using the *curves* function were subsampled and evenly spaced, resulting in 34, 34, and 27 landmarks along the dorsal, anal and caudal fins respectively (blue circles in Fig. 2). If any structures of a specimen were visibly damaged, those structures were not digitized or included in subsequent analysis. A few dorsal and anal fins (n = 4 and 7 respectively) in otherwise good condition were positioned inconsistently relative to their proximal bases in the photographs. In order to include these fins in subsequent analyses, we developed and utilized a Mac application *FinRotate* capable of adjusting the positions of median dorsal and anal fin landmarks relative to their static proximal edges. We ensured that the area of each fin changed less than 6.5% during these adjustments. These fins are indicated in table S2 and the details of each fin adjustment are given in table S3. Most balistoid fishes possess a mobile ventral keel that can be extended and retracted. For photographs in which the ventral keel was not positioned in its fully extended position (indicated with an asterisk in table S2), the *estimate.missing* function in the R package *geomorph* was used to estimate the position of the pelvic fin rudiment at the ventral tip of the extended ventral keel (Adams et al., 2017). The missing landmark positions were estimated for individuals of each species separately using all correctly positioned individuals of the same species as reference specimens. As all *C. macrocerus* individuals within the size range of the those included in the swimming trials were formalin fixed with the ventral keel depressed, a photograph of one slightly smaller, but correctly positioned, *C. macrocerus* individual was identified in the literature (Randall, 1964) and used to estimate the position of the extended ventral keel for this species. This reference specimen was not used in subsequent morphometric analyses.

**Fig. 2:**
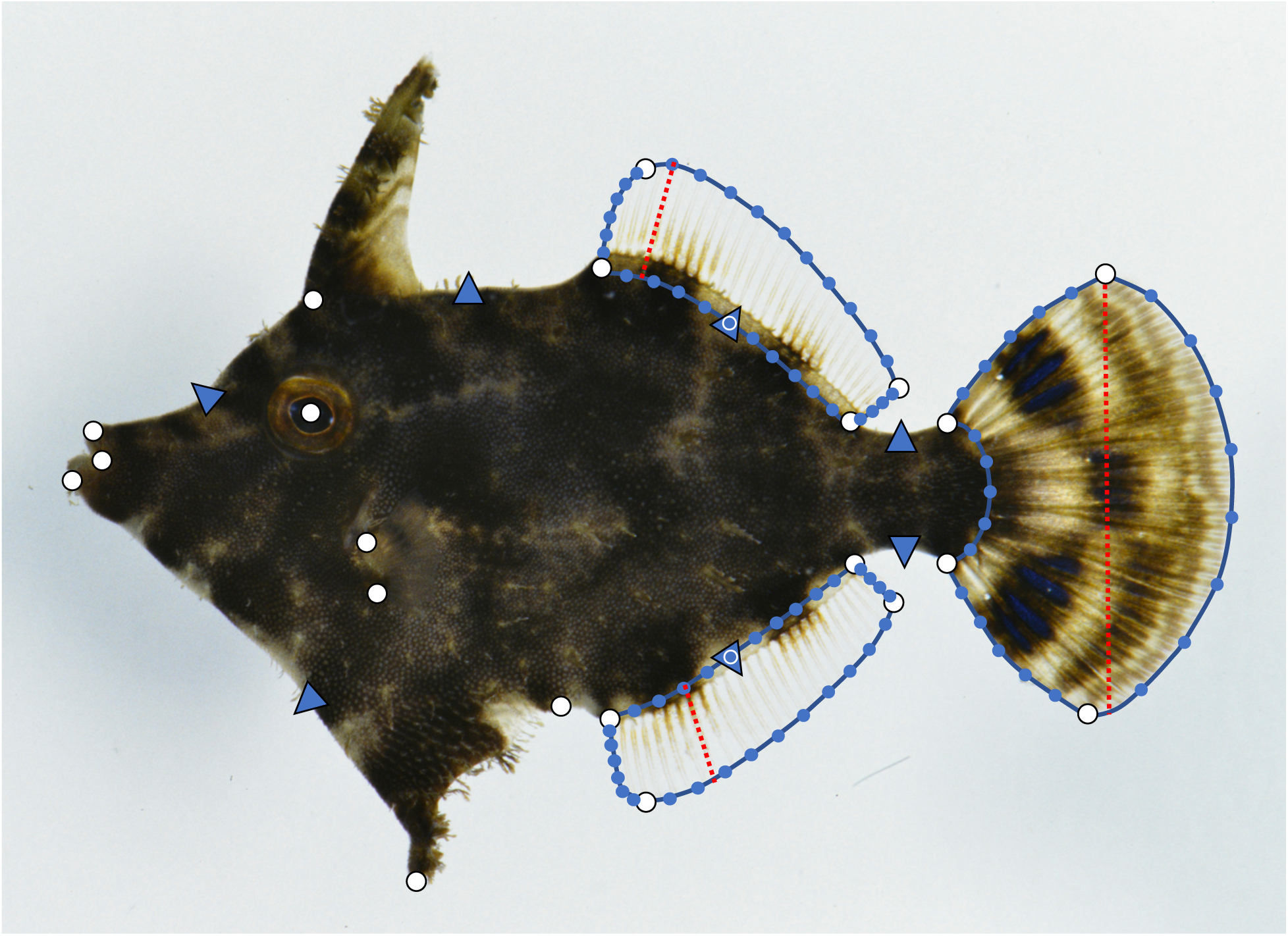
Geometrie morphometrics digitization scheme demonstrated on the filefish *Acreichthys tomentosus*, KPM-NR 53324. White circles represent functionally homologous non-sliding landmarks. Blue triangles represent sliding semi-landmarks included in only the *full shape* and *body only* datasets. Blue curves outlining the dorsal, anal and caudal fins were used for area calculations, and points along these curves (blue circles) were subsampled and treated as sliding semi-landmarks in geometric morphometric analyses. Red dotted lines represent the span measurements used for aspect ratio calculations. (photo: Hiroshi Senou).

The digitized landmark data were subdivided into five separate datasets for geometric morphometric analyses: full shape (all landmarks), body only, dorsal fin only, anal fin only, and caudal fin only. Next, each landmark was designated as either a functionally homologous *landmark* (white shapes in Fig 2) or a geometrically-relative *semilandmark* (blue shapes in Fig 2). The landmarks and semilandmarks included in each morphological dataset were then projected into tangent space using generalized Procrustes analysis (GPA) in order to remove variation in landmark position due only to rotation, translation and scaling using the *gpagen* function in the R package *geomorph* (Adams et al., 2017). During Procrustes superimposition, each *semilandmark* was allowed to slide relative to neighboring static *landmarks* using a method that minimizes the bending energy between specimens. In addition to translating, rotating and scaling the relative landmark positions of each specimen while preserving the important shape information, the *gpagen* function also calculates the centroid size (a measure of relative size) for each specimen. Each morphological dataset (full shape, body, dorsal fin, anal fin, and caudal fin) underwent GPA independently, so that Procrustes-transformed shapes and centroid sizes of each morphological unit could be analyzed separately.

Following GPA, we calculated species means of Procrustes-transformed landmark coordinates and centroid sizes for each morphological dataset. Principal components analyses (PCA) were then performed on the species-averaged, Procrustes-aligned landmark coordinates of each morphological dataset in order to identify, describe and quantify major axes of shape variation within the 13 species examined in this study using the *PlotTangentSpace* function in the R package *geomorph* (Adams et al., 2017).

The shape changes described by the two most significant axes of morphological variation (PCs 1 and 2) of each dataset were then visualized using *backtransform morphospaces* (MacLeod, 2009; Olsen, 2017). Performing PCA on the species-averaged geometric morphometric data yields a matrix of eigenvectors and PC scores for each species. Multiplying the full matrix of PC scores by the inverse of the eigenvector matrix results in a matrix of the original landmark coordinates. Trimming the PC score matrix to include only the scores along the axes of interest (PCs 1 and 2 in our case), allows one to utilize this matrix multiplication procedure to calculate theoretical landmark coordinates composed of shape variation along *only* PC axes 1 and 2. These shapes can then be plotted into a two-dimensional morphospace in order to visualize and describe shape changes along each PC axis (Olsen, 2017). We used this *backtransformation* method on the first two PC scores of each morphometric dataset in order to visualize the theoretical fin and body shapes of each species as described *only* by morphological characteristics relevant to the two most significant axes of shape variation. In order to visualize the full range of theoretical shapes in this PC1-PC2 morphospace, we generated evenly-spaced PC scores along the observed ranges of the first two axes of shape variation (PCs 1 and 2) for each dataset. We then constructed a matrix using these evenly-spaced PC scores, and multiplied this PC score matrix by the inverse of the original eigenvector matrix for each dataset. Next, we plotted the resultant theoretical shapes in their respective positions in the morphospace bounded by PCs 1 and 2 for visualization. Finally, we projected a phylogeny (McCord and Westneat 2016) into this morphospace using the *phylomorphospace* function in the R package *phytools* (Revell, 2012) in order to assess evolutionary directionality of shape changes within and between each family.

In addition to the geometric-morphometric datasets described above, five morphological ratios were calculated for each species. First, we calculated the aspect ratios (AR) of the dorsal, anal and caudal fins. Aspect ratio was calculated as,

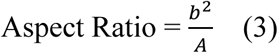

where *b* is the maximum span of the fin perpendicular to the direction of forward motion (dotted red lines in Fig. 2) and *A* is the surface area of the fin. For the dorsal and anal fins, span was measured as the Euclidian distance from base to tip of the longest fin ray of each fin independently. Caudal fin span was measured as the Euclidian distance from the dorsal-most point of the fin to the ventral-most point of the fin when the caudal fin was fully spread open. Span and area measurements were calculated using the photos digitized in *StereoMorph*. Finally, in order to assess variation in relative fin sizes between species, we calculated two area ratios. The first area ratio (*Median Fins: BCF Area Ratio*) provides a measure of relative combined dorsal and anal fin area compared to the combined body and caudal fin area. The second area ratio (*Caudal: Body Area*) provides a measure of relative caudal fin size.

### Statistical analyses

All statistical analyses were carried out in a phylogenetic context, using a pruned time-calibrated phylogeny based on 4 mitochondrial and 5 nuclear gene sequences from 80 balistoid species and 6 outgroup taxa from the families *Diodontidae, Ostraciidae* and *Tetraodontidae* (McCord and Westneat, 2016). In order to account for numerical imprecision due to small rounding errors when reading this phylogeny into R, we used the *nnls* method in the *force.ultrametric* function within the *phytools* R package (Revell, 2012) to force the tree to conform to the strict ultrametric requirements of R packages used in downstream analyses. In order to assess relationships between fish size and size-adjusted swimming metrics (in SL s^-1^), univariate phylogenetic generalized least squared (PGLS) regressions were conducted between species-averaged fish standard lengths and species-averaged, size-adjusted swimming performance metrics (U_crit_, U_t low_, U_t high_, and percent balistiform locomotion) using the *pgls* function in the *caper* R package (Orme, 2013). PGLS regressions were also conducted between all swimming performance metrics in order to identify correlations between these swimming metrics. PGLS regressions between species-averaged centroid sizes and GPA-transformed coordinates of each geometric-morphologic dataset were conducted in order to assess lingering allometric relationships between size and shape of each structure following GPA. Only the anal fin dataset exhibited a significant allometric correlation between centroid size and GPA-transformed shape (PGLS: p = 0.015), so centroid size was used as a covariate in all subsequent geometric-morphometric functional morphology PGLS regressions for the anal fin. Linear regressions were used to assess correlations between geometric morphometric PC scores and fin aspect and area ratios.

Functional morphology hypotheses were tested using univariate PGLS regressions between the species-averaged fin ratios and PC scores of the first two axes of shape variation for each geometric-morphometric dataset versus size-adjusted, species-averaged swimming performance metrics (U_crit_, U_t low_, U_t high_, and percent balistiform locomotion). In order to account for multiple statistical tests being conducted on the same dataset, all functional morphology p-values were adjusted to control for false-discovery rate using the *Benjamini-Hochberg* (BH) method (Benjamini and Hochberg, 1995) with the function *p.adjust* in the R *stats* package. All functional morphology trends were nearly identical between U_t low_ and U_t high_ (See table S4), so U_t high_ functional morphology relationships are not discussed below or included in the BH p-adjust method, resulting in a total of 45 functional morphology statistical tests input into the BH p-adjust function. The BH-adjusted p-values for functional morphology relationships are reported in the subsequent text, and raw p-values can be found in table S4. Results were considered significant when the BH-adjusted p-values were less than 0.05. All other statistical tests (correlations between measured swimming variables, allometric relationships between size and shape, phylogenetic signal in shape data, and correlations between fin ratios and geometric morphometric PC scores) were undertaken prior to hypothesis testing in order to ensure that subsequent functional morphology analyses controlled for any confounding correlations between these variables, and thus the p-values of these tests were not adjusted using the BH method. All statistical analyses were performed using the software *R* version 3.3.2 (R Core Team, 2016).

## RESULTS

### Swimming performance

The critical swimming performance (U_crit_) of the 13 balistoid species examined ranged from 4.21 SL s^-1^ in the triggerfish *B. conspicillum* to 7.05 SL s^-1^ in the triggerfish *O. niger* with an average of 5.44 SL s^-1^ (Table S5). Five individuals (one *O. niger*, two *R. aculeatus* and two *X. auromarginatus*) reached the maximum speed of the flow tank before exhaustion, so these species’ U_crit_ values should be considered conservative estimates. The U_crit_ family averages were 5.73 and 4.97 SL s^-1^ for the triggerfishes and filefishes, respectively. The initial gait transition speed (U_t low_) ranged from 2.52 SL s^-1^ in *B. conspicillum* to 5.43 SL s^-1^ in *O. niger* with an average of 3.97 SL s^-1^ (Table S5). The U_t low_ family averages were 3.73 and 4.36 SL s^-1^ for the triggerfishes and filefishes, respectively. Two filefishes, one *P. prionurus* and one *O. longirostris*, never fully transitioned to the unsteady BCF gait (U_t high_). Among fishes that did make the second gait transition, U_t high_ ranged from 3.34 SL s^-1^ in the triggerfish *R. aculeatus* to 6.12 SL s^-1^ in *O. niger* (Table S5). The U_t high_ family averages were 4.46 SL s^-1^ and 5.07 SL s^-1^ for the triggerfishes and filefishes, respectively. The percentage of the critical swimming trial in which the fishes swam using their median fins only (percent balistiform locomotion) ranged from 44.3% in *R. aculeatus* to 94.2% in *P. prionurus* (Table S5). All filefish species in this study used balistiform locomotion exclusively (without augmenting propulsion with the caudal fin) for greater than 75% of their critical swimming tests, with a family average of 88%. Conversely, all triggerfish species used balistiform locomotion alone for less than 78% of their swimming tests, with a family average of 64%.

PGLS regressions revealed no significant relationships between any size-adjusted (SL s^-1^) swimming performance metrics and fish SL (p > 0.3) (Table S4). Critical swimming speed was significantly correlated with both U_t low_ and U_t high_ (PGLS: p = 0.002 and 0.04, respectively), but not with percent balistiform locomotion (PGLS: p = 0.51). Percent balistiform locomotion was significantly correlated with both U_t low_ and U_t high_ (PGLS: p = 0.002 and 0.009, respectively). Finally, a PGLS regression revealed a highly significant correlation between U_t low_ and U_t high_ (p = 0.002). Nearly all trends between morphology and gait transition speeds were the same regardless of which gait transition speed was used (Table S4), so subsequent results will include only U_t low_, Ucrit, and percent balistiform locomotion.

### Fin ratios

Anal fin aspect ratios (AR) ranged from 0.32 in *O. longirostris* to 1.24 in *O. niger* with an average of 0.62. Triggerfishes tend to have higher AR anal fins (range = 0.52 - 1.24, mean = 0.73) than filefishes (range = 0.32 - 0.62, mean = 0.43). Dorsal fin AR ranged from 0.31 in *O. longirostris* to 1.16 in *O. niger* with an average of 0.60. Triggerfishes also tend to have higher AR dorsal fins (range = 0.46 - 1.16, mean = 0.72) than filefishes (min = 0.31, max = 0.53, mean = 0.41). Caudal fin AR ranged from 1.57 in *C. macrocerus* to 3.05 in *X. auromarginatus* with an average of 2.51. Triggerfishes tend to have higher AR caudal fins (range = 2.46 - 3.05, mean = 2.73) than filefishes (range = 1.57 - 2.70, mean = 2.16). The *Median Fin: BCF Area Ratio* ranged from 0.15 in *R. aculeatus* and *A. tomentosus* to 0.32 in *O. niger* with an average of 0.21. There was no difference between triggerfish and filefish family means for this metric. The *Caudal: Body Area Ratio* ranged from 0.13 in *B. conspicillum* to 0.32 in *P. prionurus* with an average of 0.18. Filefishes tend to have larger caudal fins relative to their bodies (range = 0.20 - 0.32, mean = 0.25) than triggerfishes (range = 0.13 - 0.16, mean = 0.14).

### Geometric morphometrics

#### Full shape

The primary axis of variation (PC1: 46%) for this dataset, including all parts of the fish, describes the length and position of the dorsal and anal fins and the caudal peduncle depth compared to body depth. High PC1 scores are associated with narrow caudal peduncles, deep bodies, and short, posteriorly positioned dorsal and anal fins. The second major axis of variation (PC2: 24%) differentiates fishes primarily based on the shape of their fins. High PC2 morphospace is occupied by fishes with concave or truncate caudal fins and high aspect ratio (AR) median fins with long leading edges and short trailing edges. Conversely, low PC2 morphospace is occupied by fishes with highly convex caudal fins and low AR median fins with leading and trailing edges of similar lengths. Filefishes cluster in the area of full shape morphospace defined by shallow bodies, wide caudal peduncles and long median fins (low PC1) as well as convex caudal fins and low AR median fins (low PC2). Triggerfishes occupy the area of morphospace defined by deep bodies, narrow caudal peduncles and short median fins (high PC1) as well as convex or truncate caudal fins and mid to low AR median fins (low to mid PC2), with the exception of *O. niger*, which sits in a largely unoccupied area of morphospace defined by low PC1 and high PC2 scores (Fig. 3A).

**Fig. 3:**
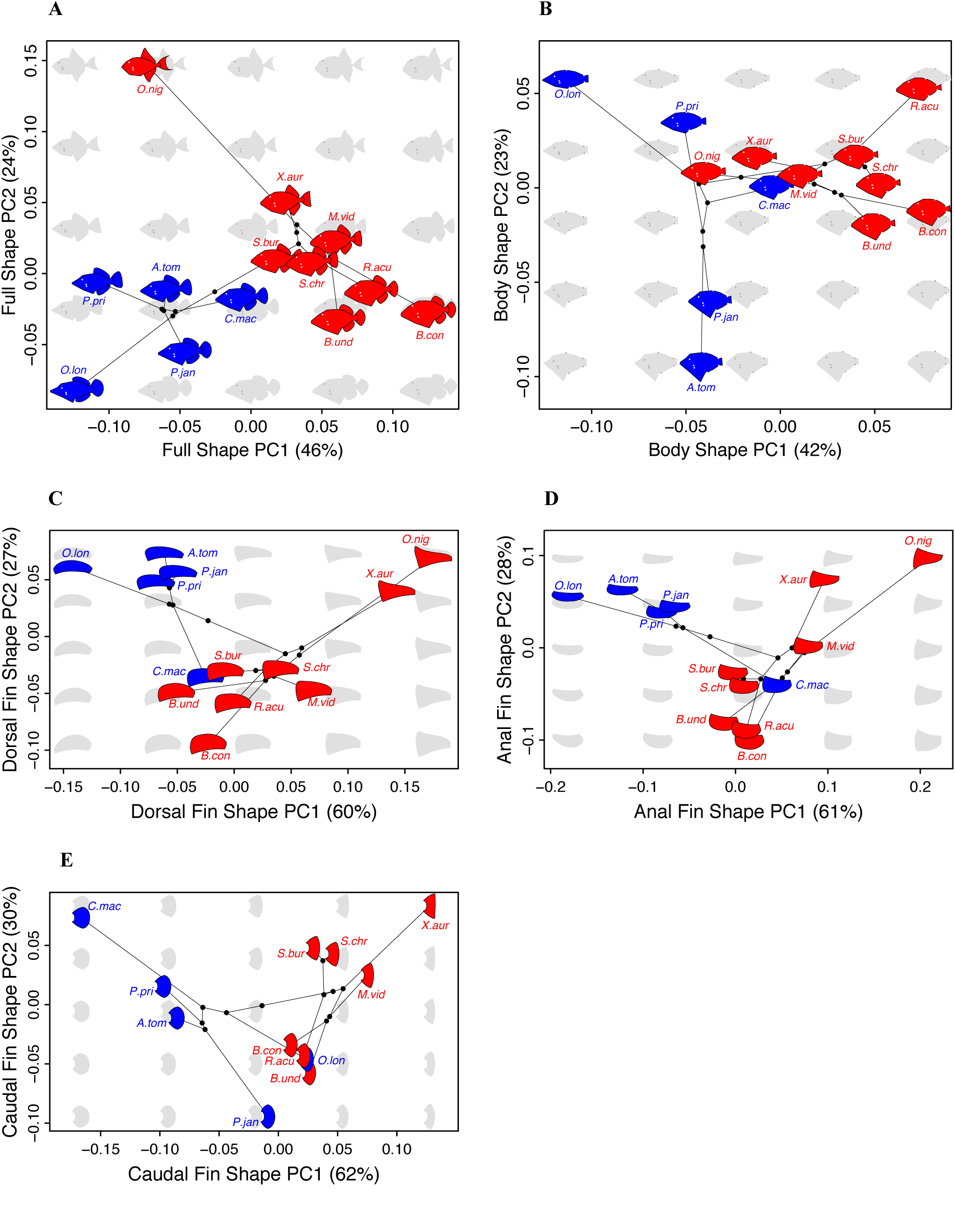
Backtransformation phylomorphospaces depicting theoretical shapes along the two most significant axes of shape variation for each morphological dataset. Red (Balistidae) and blue (Monacanthidae) shapes represent the backtransform shapes of each species included in this study. Gray shapes represent theoretical backtransform shapes corresponding to each location in morphospace. Black lines represent the phylogeny from Fig. 1 transformed into each morphospace. The black dots between these lines represent theoretical positions of each ancestral node. (A) Full shape. (B) Body only. (C) Dorsal fin only. (D) Anal fin only. (E) Caudal fin only. Samples sizes for each morphometric dataset reported in table S1.

#### Body only

The primary axis of variation (PC1: 42%) describes the ratio of anterior body depth to posterior body depth, as well as slope of the head profile and median fin length. Fishes with high PC1 values are deep-bodied anteriorly with narrow caudal peduncles, short median fins, and convex forehead profiles. Conversely, fishes occupying areas of low PC1 morphospace are shallow-bodied anteriorly with wide caudal peduncles, long median fins and a concave forehead profile. Principal component 2 (PC2: 23%) describes the depth of the ventral keel. Fishes with deep ventral keels occupy low areas of PC2 morphospace. Most triggerfishes cluster in areas of morphospace defined by deep bodies, narrow caudal peduncles, convex forehead profiles (mid to high PC1) and ventral keels of moderate to shallow depth (mid to high PC2). Filefishes generally occupy areas of morphospace defined by narrow bodies, wide caudal peduncles and concave forehead profiles (mid to low PC1) and span the entire range of PC2 morphospace. The filefish *C. macrocerus* appears to have converged upon an area of body morphospace primarily occupied by triggerfishes (Fig. 3B).

#### Dorsal fin

The primary axis of variation (PC1: 60%) describes the length ratio of the leading edge to the trailing edge of the fin. Low PC1 regions of morphospace are occupied by fins with leading and trailing edges of similar lengths, while high PC1 regions are occupied by fins with long leading edges and short trailing edges. Increasing dorsal fin PC1 scores are associated with increasing dorsal fin aspect ratios (AR) (linear regression: p < 0.001, R^2^ = 0.96). Principal component 2 (PC2: 27%) differentiates dorsal fins based on overall length-depth ratio, with long and shallow fins occupying areas of higher PC2 morphospace, and short, deep fins occupying areas of low PC2 morphospace. Dorsal fin PC2 is not associated with fin AR (linear regression: p = 0.87, R^2^ = 0.002). Filefishes cluster in the top left corner of morphospace (low PC1, high PC2) defined by low AR, shallow and elongate fins with the exception of *C. macrocerus*, which groups with the majority of the triggerfishes in the area of morphospace defined by short, deep dorsal fins of intermediate AR (mid PC1, low PC2). *Odonus niger* and *X. auromarginatus* have diverged from the rest of the Balistidae to occupy the top right corner of morphospace (high PC1 and high PC2) defined by high AR, elongate dorsal fins (Fig. 3C).

#### Anal fin

The primary axis of morphological variation (PC1: 61%) describes the overall length-depth ratio of the fin as well as the orientation and length ratio of leading edge to trailing edge of the fin. Low PC1 regions of morphospace are occupied by elongate, shallow anal fins, with leading and trailing edges of nearly equal lengths. Conversely, high PC1 regions of morphospace are occupied by short, deep anal fins with anteriorly-oriented leading edges significantly longer than their trailing edges. Increasing anal fin PC1 values are associated with increasing anal fin aspect ratios (AR) (linear regression: p < 0.001, R^2^ = 0.90). Principal component 2 (PC2: 28%) differentiates anal fins primarily by the length of the leading edge and shape of the distal edge of the fin. Anal fins with high PC2 scores have anteriorly oriented, straight leading edges with posteriorly tapering distal edges, while fins with low PC2 scores have posteriorly-curved leading edges and deep, convexly rounded distal edges. Anal fin PC2 is not correlated with anal fin AR (linear regression: p = 0.42, R^2^ = 0.058). Filefishes cluster in morphospace defined by shallow, elongate anal fins (low PC1) with anteriorly-oriented leading edges and slightly taping distal edges (high PC2), with the exception of *C. macrocerus*, which groups with the majority of the triggerfishes in the area of morphospace defined by deep, rounded anal fins (low PC2) of intermediate length and AR (mid PC1). Once again, *O. niger and X. auromarginatus* fall near the top right corner (high PC1 and high PC2) defined by elongate, posteriorly tapering, high AR anal fins (Fig. 3D).

#### Caudal fin

The forked caudal fin shape of *O. niger* is a major outlier along PC1 within the caudal fin geometric-morphometric dataset according to a Rosner’s generalized extreme Studentized deviate test conducted with the *rosnerTest* function in R (Millard, 2013), so this species was removed from all caudal fin geometric-morphometric analyses and the caudal fin PCA was rerun without *O. niger*. Among the remaining 12 species, the primary axis of variation (PC1: 62%) describes changes in overall length-depth ratios and the shape of the posterior edge of the caudal fin. Caudal fins occupying areas of low PC1 morphospace are elongate and narrow with highly convex posterior edges, while caudal fins occupying areas of high PC1 morphospace are short and deep with truncate posterior edges. Principal component 2 (PC2: 30%) is highly correlated with caudal fin aspect ratio (AR) (linear regression: p < 0.001; R^2^ = 0.91). High PC2 areas of morphospace are occupied by high AR caudal fins, while low PC2 areas are occupied by low AR caudal fins. Triggerfishes possess fairly wide caudal fins with slightly convex or truncate distal edges (mid to high PC1). Most filefishes possess fairly narrow caudal fins with more convex distal edges (mid to low PC1). Fishes from both families span the entirety of PC2 (Fig. 3E).

### Functional morphology

#### Fin ratios and performance

Univariate PGLS regressions revealed significant positive correlations between U_crit_ and dorsal, anal and caudal fin aspect ratios (AR) (see table 1 for statistical significance) (Fig. 4A-C). Additionally, increasing the area ratio of the median fins to the body and caudal fins (*Median Fins: BCF Area Ratio*) is associated with increased U_crit_ (PGLS: p = 0.0385) (Fig. 4D). Higher dorsal and anal fin ARs and increased *Median Fins: BCF Area Ratio* are also associated with increased gait transition speed (U_t low_) (Table 1) (Fig. 5). Caudal fin AR is not significantly associated with U_t low_ among balistoid fishes as a whole (PGLS: p = 0.0588), but caudal fin AR is positively correlated with U_t low_ among triggerfishes (Balistidae) alone (PGLS: p = 0.0386) (Fig. 5C). Finally, *Median Fins: BCF Area Ratio* is positively correlated with percent balistiform locomotion (PGLS: p = 0.0217) (Fig. 6).

**Table 1:** **Functional morphology results**

|  |  | Benjamini and Hochberg (BH) Adjusted P Value |  |  |
| --- | --- | --- | --- | --- |
| PGLS Relationship | Directionality | Balistoidea | Balistidae | Monacanthidae |
| Dorsal AR vs $U_{crit}$ | Pos | **0.0169** | 0.0844 | 0.965 |
| Anal AR vs $U_{crit}$ | Pos | **0.0169** | 0.0665 | 0.965 |
| Caudal AR vs $U_{crit}$ | Pos | **0.0385** | **0.0491** | 0.965 |
| Median Fins: BCF area ratio vs $U_{crit}$ | Pos | **0.0385** | 0.1098 | 0.965 |
| Dorsal AR vs $U_{t low}$ | Pos | **0.0111** | **0.0333** | 0.965 |
| Anal AR vs $U_{t low}$ | Pos | **0.0104** | **0.0333** | 0.965 |
| Caudal AR vs $U_{t low}$ | Pos | 0.0588 | **0.0386** | 0.965 |
| Median Fins: BCF area ratio vs $U_{t low}$ | Pos | **0.0017** | **0.0262** | 0.965 |
| Median Fins: BCF area ratio vs % Bal | Pos | **0.0217** | 0.0844 | 0.614 |
| Full Shape PC2 vs $U_{crit}$ | Pos | **0.0385** | 0.0954 | 0.614 |
| Body PC1 vs $U_{crit}$ | Neg | **0.0474** | 0.0844 | 0.965 |
| Dorsal PC1 vs $U_{crit}$ | Pos | **0.0217** | 0.0844 | 0.965 |
| Dorsal PC2 vs $U_{crit}$ | Pos | **0.0169** | **0.0491** | 0.965 |
| Full Shape PC1 vs $U_{t low}$ | Neg | **0.0104** | **0.0333** | 0.965 |
| Full Shape PC2 vs $U_{t low}$ | Pos | **0.0169** | **0.0333** | 0.965 |
| Body PC1 vs $U_{t low}$ | Neg | **0.0081** | **0.0262** | 0.965 |
| Dorsal PC1 vs $U_{t low}$ | Pos | **0.0169** | **0.0333** | 0.965 |
| Dorsal PC2 vs $U_{t low}$ | Pos | **0.0081** | **0.0262** | 0.965 |
| Anal PC2 vs $U_{t low}$ | Pos | **0.0219** | 0.0701 | 0.965 |
| Full Shape PC1 vs % Bal | Neg | **0.0385** | 0.139 | 0.965 |
| Body PC1 vs % Bal | Neg | **0.0385** | 0.113 | 0.965 |

PGLS = Phylogenetic generalized least squared regression. AR = aspect ratio, % Bal = percent balistiform locomotion. Bold values indicate statistically significant trends ( $p < 0.05$ ).

**Fig. 4:**
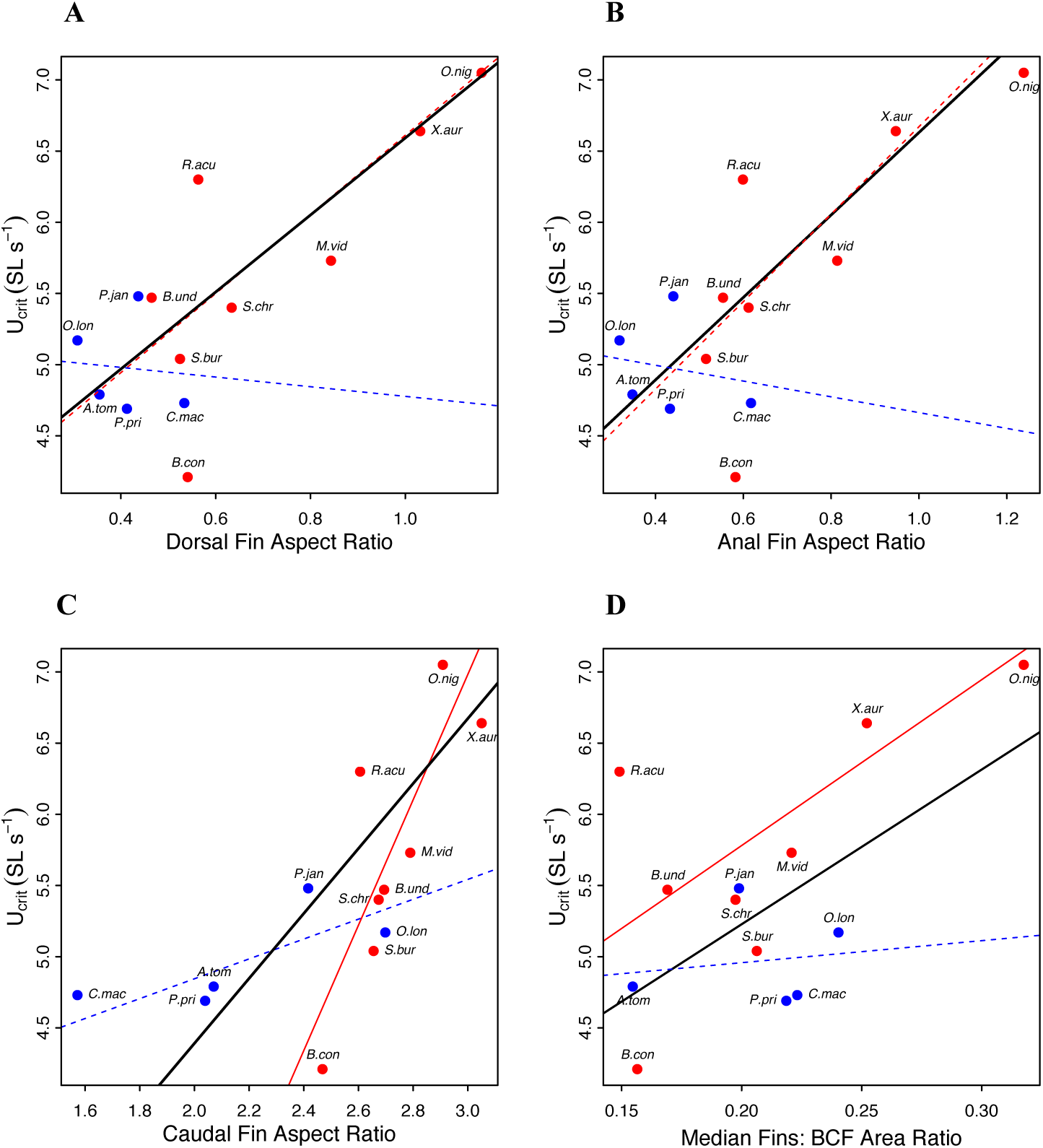
Significant trends between fin ratios and critical swimming performance (U_crit_) measured in standard lengths per second (SL s^-1^). (A) Dorsal fin aspect ratio. (B) Anal fin aspect ratio. (C) Caudal fin aspect ratio. (D) Ratio of median fin area to combined body and caudal fin area (*Median Fins: BCF Area Ratio*). Red and blue dots indicate triggerfishes and filefishes, respectively. Black lines depict the phylogenetic generalized least squared (PGLS) regression lines for all 13 balistoid species included in this study. Red and blue lines depict independent PGLS regression lines for the triggerfishes and filefishes, respectively. Solid and dotted lines indicate significant (p < 0.05) and nonsignificant (p > 0.05) trends, respectively. Samples sizes for U_crit_ and morphometric datasets are reported in tables S5 and S1, respectively.

**Fig. 5:**
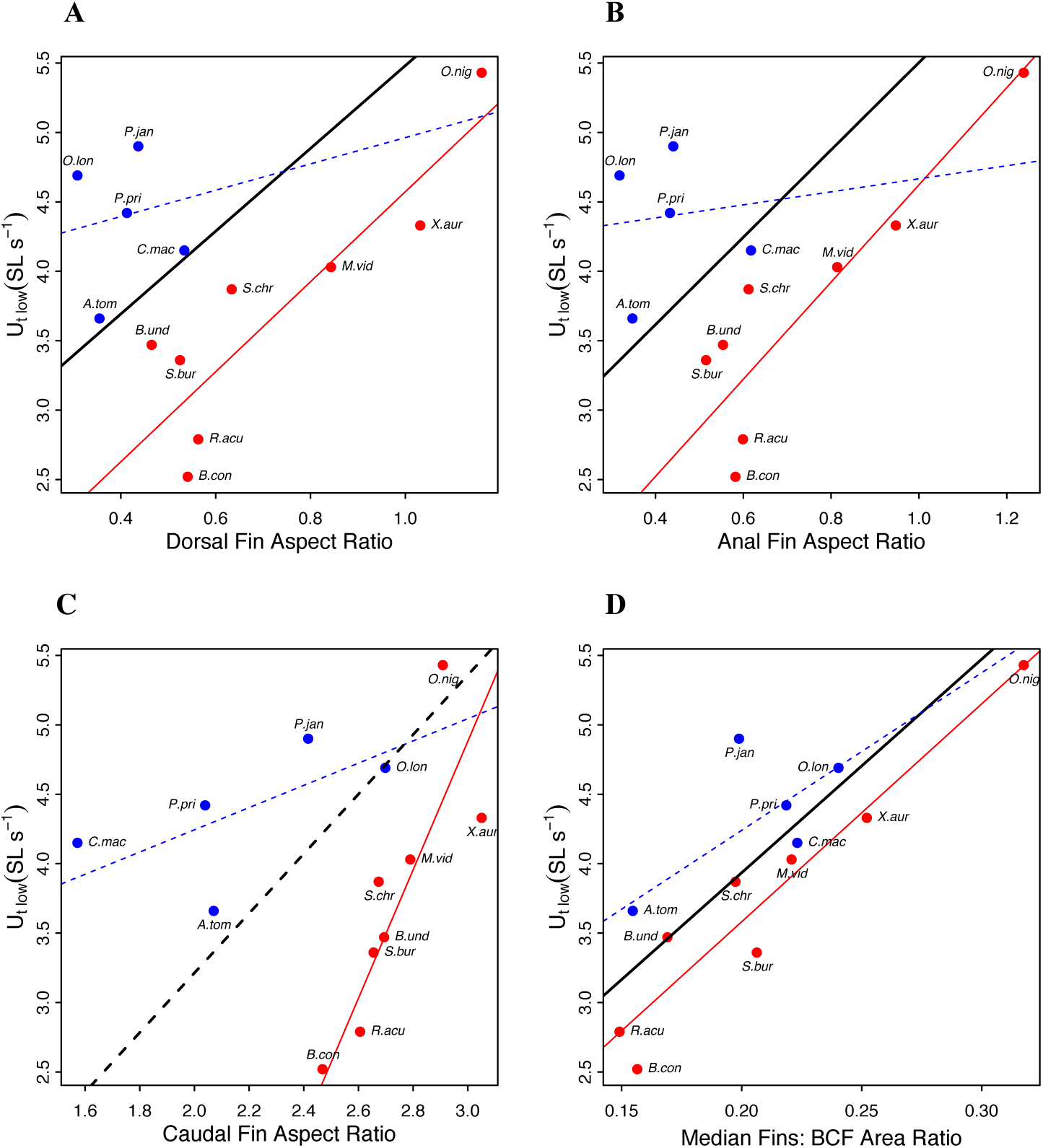
Significant trends between fin ratios and the first gait transition speed (U_t low_) measured in standard lengths per second (SL s^-1^). (A) Dorsal fin aspect ratio. (B) Anal fin aspect ratio. (C) Caudal fin aspect ratio. (D) Ratio of median fin area to combined body and caudal fin area (*Median Fins: BCF Area Ratio*). Family association and statistical significance are indicated by color and line type, respectively, as in Fig. 4. Samples sizes for U_t low_ and morphometric datasets are reported in tables S5 and S1, respectively.

**Fig. 6:**
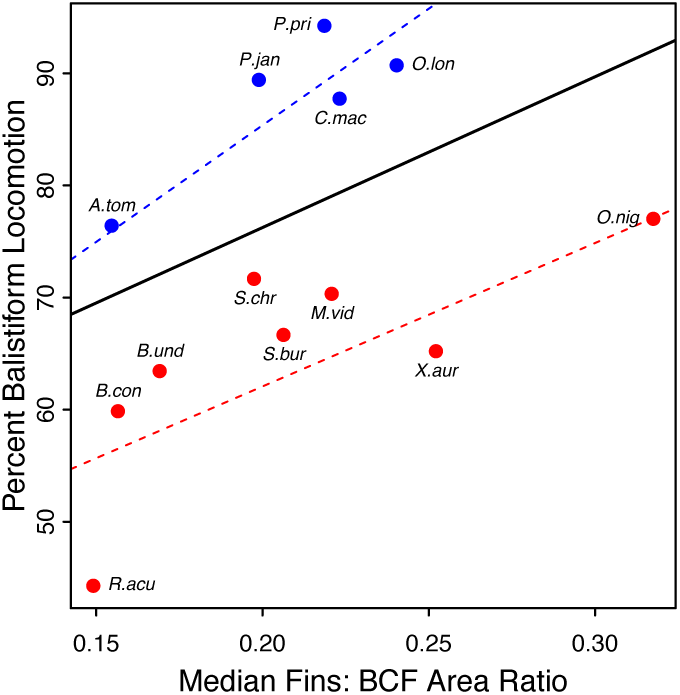
Relationship between relative median fin size and the percentage of the critical swimming speed achieved using the balistiform gait alone (percent balistiform locomotion). This scatterplot depicts the relationship between the ratio of median fin area to the combined body and caudal fin area (*Median Fins: BCF Area Ratio*) and percent balistiform locomotion. Family association and statistical significance are indicated by color and line type, respectively, as in Fig. 4. Samples sizes for percent balistiform locomotion and morphometric datasets are reported in tables S5 and S1, respectively.

#### Geometric morphometrics and performance

Univariate PGLS regressions of PC scores against swimming performance metrics, revealed a total of 12 significant functional morphology trends within the superfamily Balistoidea (Table 1). Fishes with deeper bodies and less convex caudal fins (high full shape PC2) and fishes with elongate (high dorsal fin PC2), posteriorly-tapering (high dorsal fin PC1) dorsal fins are associated with increased U_crit_ (Fig. 7). Many aspects of balistoid median fin and body shape are associated with U_t low_ (Fig. 8) (Table 1). Interestingly, the most significant axes of median fin shape correlated with U_t low_ are not correlated with fin aspect ratio (AR). Specifically, elongate dorsal and anal fins, regardless of AR (high dorsal and anal fin PC2s) are associated with higher U_t low_ (Fig. 8D,E). Although caudal fin shape alone is not associated with U_t low_, balistoid fishes with narrow caudal peduncles (high full shape PC1) *and* highly convex caudal fins (low full shape PC2) tend to recruit the caudal fin at slower speeds (lower U_t low_) than fishes with wide caudal peduncles *and* less convex caudal fins (Fig. 8B). Finally, fishes with elongate median fins *and* wide caudal peduncles (low full shape and body PC1s) use balistiform locomotion alone for a higher percentage of their overall critical swimming speed (high percent balistiform locomotion) than fishes with short median fins *and* narrow caudal peduncles (high full shape and body PC1s)(Fig. 9).

**Fig. 7:**
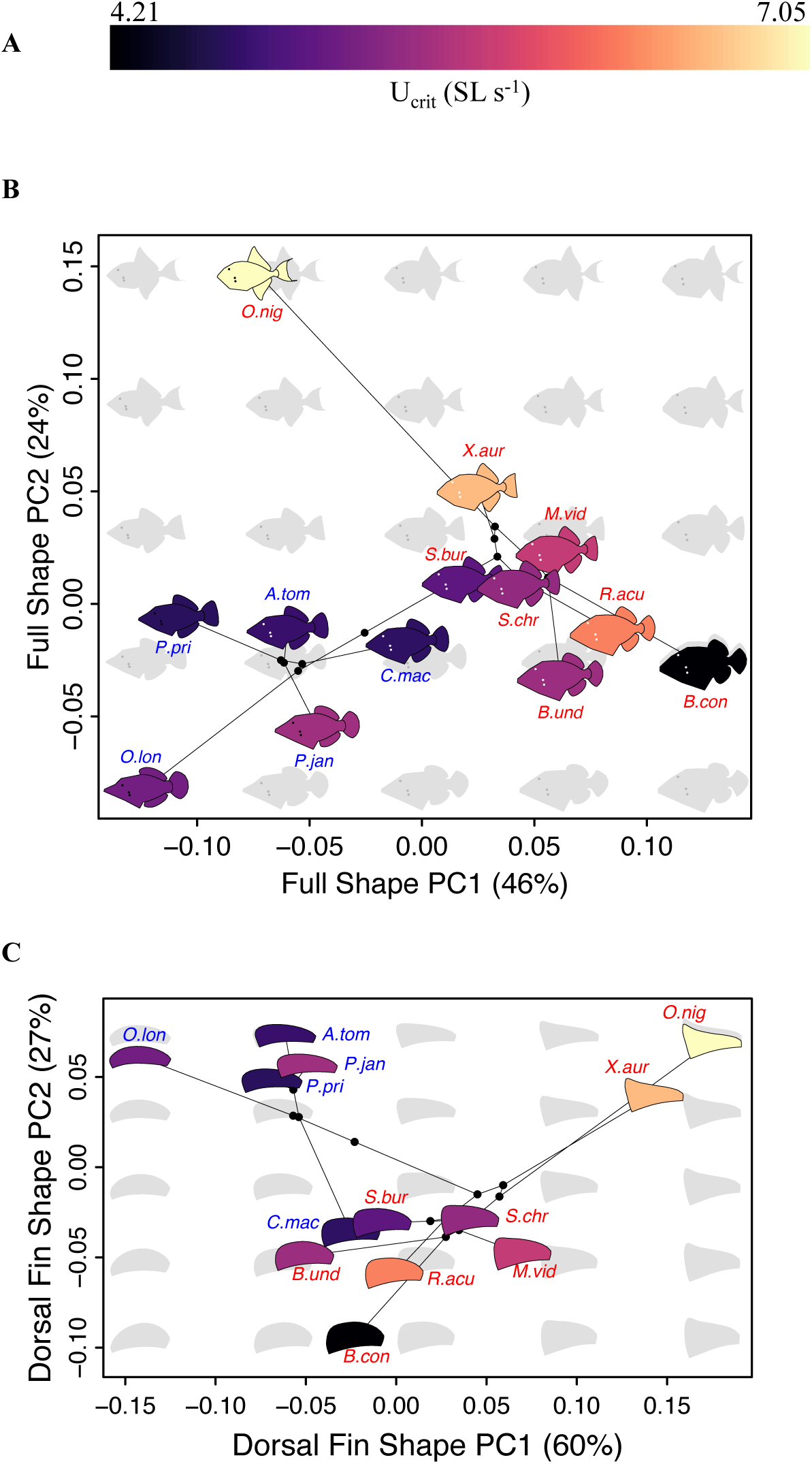
Backtransformation phylomorphospace plots color-coded for critical swimming speed (U_crit_). See Fig. 3 for description of backtransformation phylomorphospace plots. Shapes affiliated with each species have been color-coded according to U_crit_ measured in standard lengths per second (SL s^-1^). (A) The color gradient used to depict U_crit_. (B) Full shape. (C) Dorsal fin only. Species abbreviations are color-coded based on family, with triggerfishes in red and filefishes in blue. Samples sizes for U_crit_ and morphometric datasets are reported in tables S5 and S1, respectively.

**Fig. 8:**
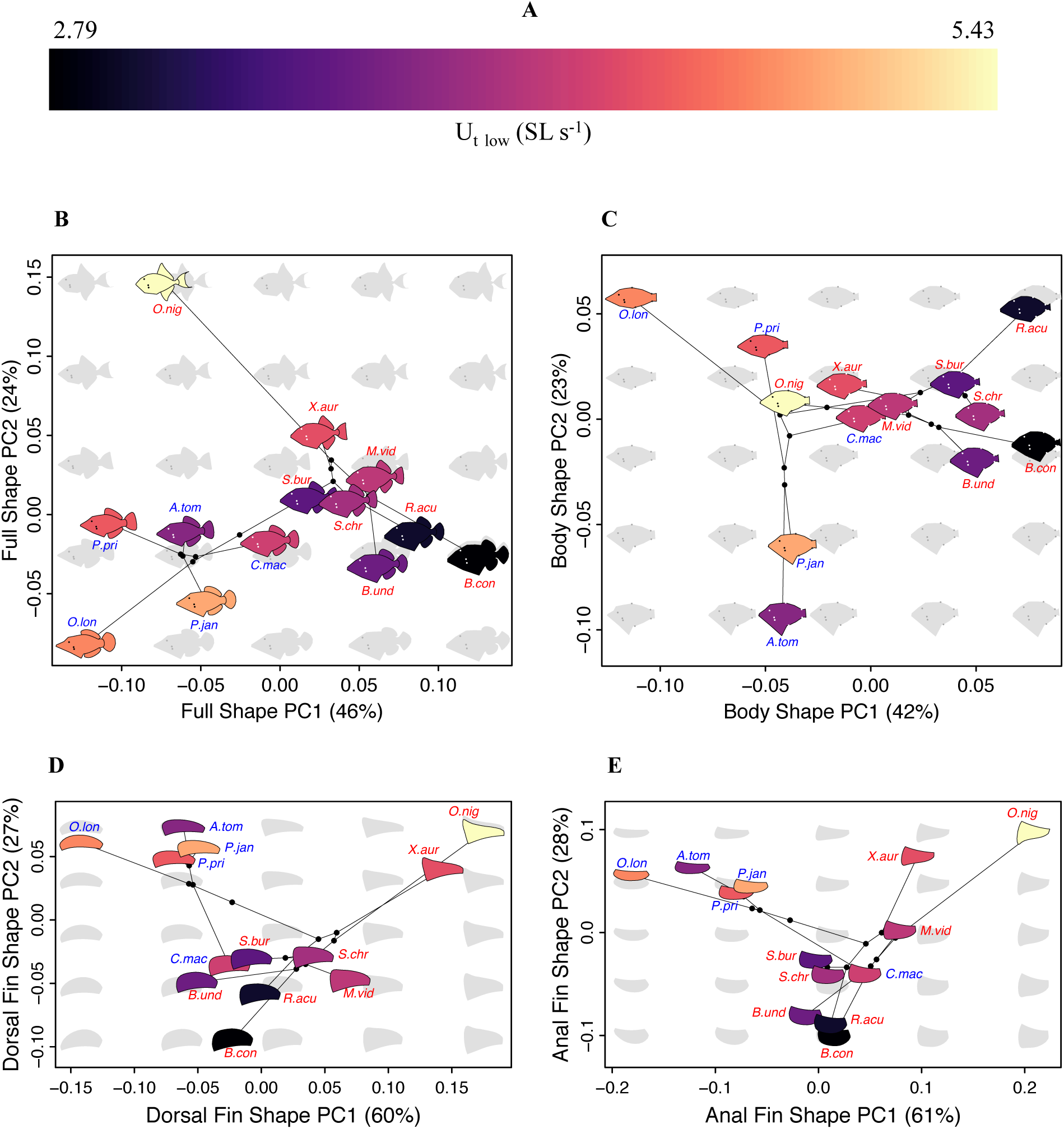
Backtransformation phylomorphospace plots color-coded for the first gait transition speed (U_t low_). See Fig. 3 for description of backtransformation phylomorphospace plots. Shapes affiliated with each species have been color-coded according to U_t low_ measured in standard lengths per second (SL sec^-1^). (A) The color gradient used to depict the speed at which the first gait transition (U_t low_) occurred. (B) Full shape. (C) Body only. (D) Dorsal fin only. (E) Anal fin only. Species abbreviations are color-coded based on family, with triggerfishes in red and filefishes in blue. Samples sizes for U_t low_ and morphometric datasets are reported in tables S5 and S1, respectvely.

**Fig. 9:**
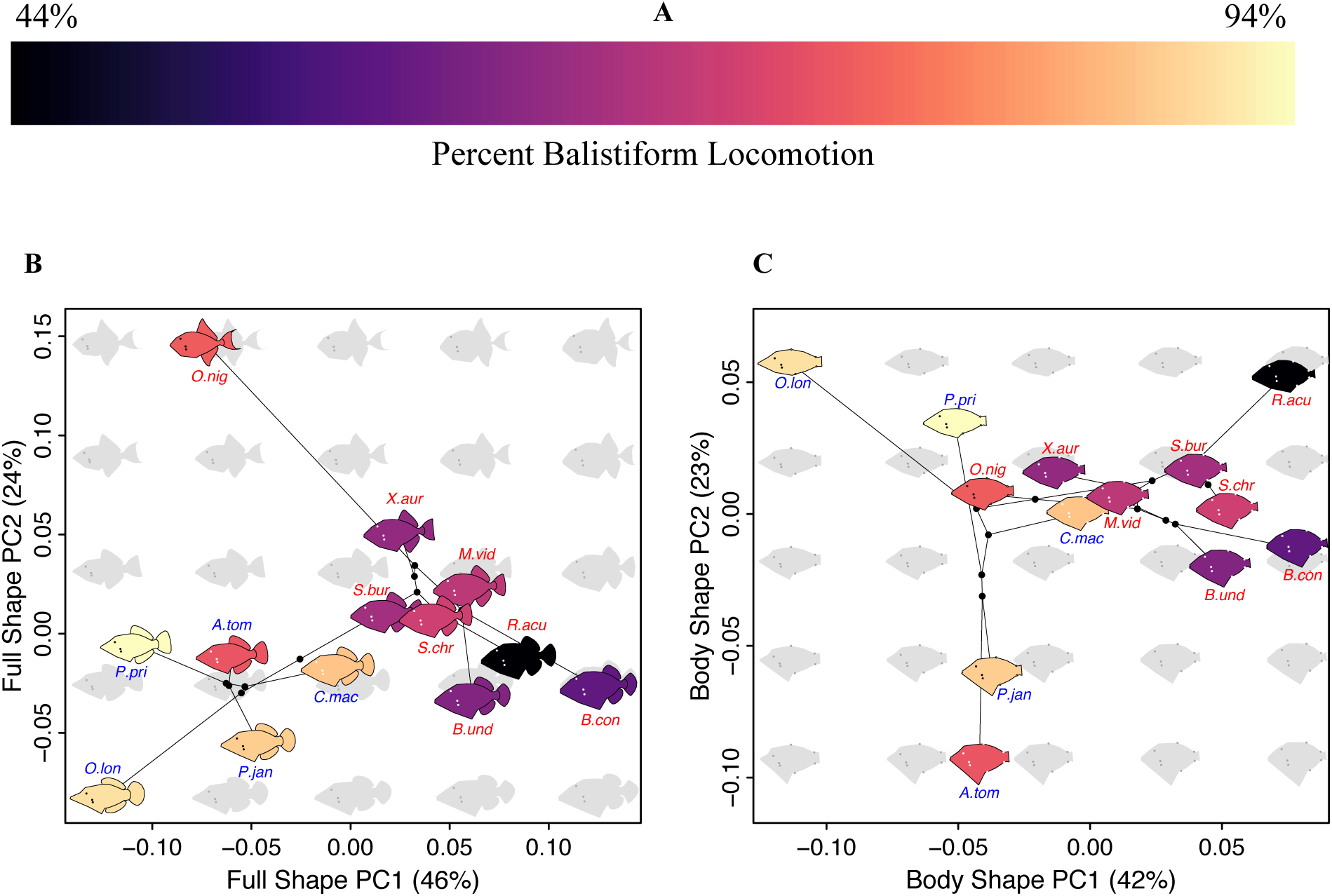
Backtransformation phylomorphospace plots color-coded for percentage of the critical swimming speed achieved using the balistiform gait alone (percent balistiform locomotion). See Fig. 3 for description of backtransformation phylomorphospace plots. Shapes affiliated with each species have been color-coded according to percent balistiform locomotion. (A) The color gradient used to depict percent balistiform locomotion. (B) Full shape. (C) Body only. Species abbreviations are color-coded based on family, with triggerfishes in red and filefishes in blue. Samples sizes for percent balistiform locomotion and morphometric datasets are reported in tables S5 and S1, respectively.

#### Family trends

Many of the trends in functional morphology remain significant when assessed within the eight triggerfish species alone, but many filefish species do not follow the overall balistoid functional morphology trends (Table 1). In fact, among the five filefish species examined in this study alone, no axes of body or fin shape variation are associated with U_crit_, U_t low_, or percent balistiform locomotion (Table 1, Table S4).

## DISCUSSION

Triggerfishes and filefishes are capable of relatively high performance steady swimming using the balistiform gait alone, comparable to median and paired-fin locomotor specialists from other fish families. The hypothesis that fin aspect ratio is associated with higher endurance swimming performance and a higher gait transition speed was strongly supported in our analysis. However, triggerfishes and filefishes showed different endurance swimming strategies and gait transition behaviors, with filefishes showing a strikingly high performance using pure balistiform locomotion. The central conclusions of this study are (1) fin and body shape are good predictors of overall critical swimming performance in balistoid fishes, (2) balistoid fishes exhibit a variety of gait transition strategies to achieve high-speed endurance swimming performance and (3) each species appears to be morphologically specialized to take advantage of one of these gait transition strategies.

### Swimming performance

The results of this study indicate that balistoid fishes are capable of strong endurance swimming performance. The fastest species in this study, *O. niger*, achieved an average critical swimming speed of 7.05 SL s^-1^, similar to fishes such as wrasses (Walker and Westneat, 1997) that are considered to be high-performance paired-fin endurance swimmers. The swimming performance results from this study also reveal interesting trends between balistoid families. Triggerfishes tend to achieve faster overall critical swimming speeds than filefishes (5.73 and 3.97 SL s^-1^, respectively). However, triggerfishes recruit body and caudal fin musculature (U_t low_) at lower speeds than filefishes (3.73 and 4.36 SL s^-1^, respectively) and thus power swimming with balistiform locomotion alone for a lower percentage of their overall endurance swimming performance than filefishes (65% and 88%, respectively). This means that, on average, filefishes actually achieved faster swimming speeds than triggerfishes using balistiform locomotion alone (higher U_t low_). Furthermore, this suggests that although nearly all past studies concerning the evolution (Dornburg et al., 2011), biomechanics (Blake, 1978; Wright, 2000; Korsmeyer et al., 2002; Hu et al., 2006; Loofbourrow, 2009), energetics (Korsmeyer et al., 2002) and performance (Wright, 2000; Korsmeyer et al., 2002) of balistiform locomotion have used triggerfishes as their balistiform swimming models, filefishes are likely a better system for studying high-speed balistiform locomotion due to their heavy reliance on the balistiform gait alone throughout much of their swimming speed range. Many past studies concerning balistiform locomotion (Korsmeyer et al., 2002; Hu et al. 2006; Loofbourrow, 2009) have focused solely on *R. aculeatus*, the species that used balistiform locomotion for the smallest percentage of its total Ucrit in this study (44%), so the biomechanical and energetic trends observed in these studies may not be broadly applicable to all balistoid fishes. Keeping this in mind, it is important to note that previous research has indicated that *R. aculeatus* actually incurs a significant energetic cost when transitioning from balistiform locomotion to a combined balistiform plus BCF gait (U_t low_), suggesting that balistoid fishes might undergo gait transitions in order to meet the power requirements of swimming faster in a highly viscous aquatic environment (Korsmeyer et al., 2002). This is in stark contrast to the gait transitions of terrestrial animals, which are typically undergone in order to maximize mechanical efficiency at each speed (reviewed in Alexander, 1989). This power-requirement, rather than mechanical efficiency, energetics pattern described in *R. aculeatus* suggests that our U_t low_ measurements may be a good estimate of the upper speed limit of the balistiform gait for each species. However, this energetics pattern might not apply to all balistoid species, especially those with very different morphologies and fin kinematics. Either way, it is clear from our results as well as those of Korsmeyer et al. (2002) that not all balistoid fishes power aerobic endurance swimming using the balistiform gait alone, but rather are capable of long-term aerobic locomotion using the combined balistiform and BCF gait.

### Functional morphology

Critical swimming performance of balistoid fishes is highly correlated with fin aspect ratios (AR) and area ratios. Fishes with high AR dorsal, anal and caudal fins and fishes with relatively large median fins tend to achieve higher Ucrit speeds. These trends support previous findings that balistoid fishes benefit from hydrodynamically efficient (high AR) and powerful (relatively large) dorsal and anal fins during endurance swimming (Wright, 2000). By expanding the scope of performance measures and fin metrics to additional species, this trend is also expanded to include filefishes. The relationship between increasing median fin ARs and increasing critical swimming performance is not surprising, given the hydrodynamic benefits of high AR fins (Lighthill, 1970; Bushnell and Moore, 1964) and the fact that the median fins are significantly involved in thrust production throughout the majority of the critical swimming tests (through U_t high_). The relationship between increasing caudal fin AR and overall critical swimming performance reflects the fact that, on average, these balistoid fishes started recruiting occasional caudal fin oscillations (U_t low_) at speeds only 74% of the way to their respective critical swimming limits. Triggerfishes were especially reliant on caudal fin contribution, with a family average percent balistiform locomotion of only 64%. Theoretical work has demonstrated many hydrodynamic advantages of high AR fins, including decreased drag and decreased production of destabilizing tip vortices (Bushnell and Moore, 1964; Lighthill, 1970; Xin and Wu, 2013). These hydrodynamic advantages likely make balistoid fishes with high AR median and caudal fins capable of more energetically efficient propulsion using the balistiform and BCF gaits, respectively.

The seven strongest trends between morphology and swimming performance measured in this study were associated with gait transition speed (U_t low_), rather than overall critical swimming performance. Specifically, fishes with large median fins, high aspect ratio (AR) median fins, long median fins (regardless of AR), and wide caudal peduncles recruited axial musculature and the caudal fin (U_t low_) at higher speeds than fishes with small, short and rounded median fins and narrow caudal peduncles. It is important to point out that the filefishes in this study are outliers in the trend between increasing median fin aspect ratio and increasing U_t low_. In fact, close examination of these trends (Fig. 5A,B; Fig. 8D,E) actually suggests that possessing a median fin shape at either extreme of the AR continuum or median fin PC1 range (significantly associated with AR) results in improved swimming performance while using the balistiform gait alone (U_t low_) compared to fishes with median fins of intermediate ARs. Specifically, the species with the second, third and fourth fastest U_t low_ speeds (*P. janthinosoma, O. longirostris*, and *P. prionurus*) actually possess some of the lowest AR median fins measured in this study. In other words, it appears that two optima may exist between median fin AR and swimming performance powered by the balistiform gait alone (U_t low_), with the fastest balistiform locomotion speeds (U_t low_) achieved by fishes with the highest *and* lowest AR median fins, and the slowest maximum balistiform swimming speeds (U_t low_) achieved by fishes with median fins of intermediate ARs.

The multiple optima discovered between median fin aspect ratio (AR) and U_t low_ can likely be explained by the fin kinematics used to achieve these fast sustained balistiform locomotion speeds. Kinematics research has shown that high AR median fin are associated with highly oscillatory, flapping balistiform fin kinematics, while low AR median fins are associated with more wave-like, undulatory median fin kinematics (Wright, 2000). All the filefishes added to this study (*A. tomentosus, O. longirostris, P. prionurus* and *P. janthinosoma*) have lower AR median fins than the fishes included in the Wright (2000) study, suggesting that these filefishes may possess even more undulatory fin kinematics than previously described for the group, although fin kinematics must be experimentally confirmed. The differences in median fin kinematics used by the balistiform swimmers at either end of the median fin AR spectrum likely place different hydrodynamic pressures on the fishes (Wright, 2000; Sprinkle et al. 2017), and each fin shape may be hydrodynamically optimized for its respective kinematics. It is important to note that although we see multiple optima in the relationship between median fin AR and U_t low_, we do see a strong linear relationship between increasing median fin AR and increasing U_crit_, suggesting that fishes with short, intermediate AR median fins with low U_t low_ performance compensate with high-performance BCF gaits.

Associations between body and caudal fin shape and gait transition speed can also be explained by hydrodynamic principals. We found that balistoid fishes with bodies that taper posteriorly into narrow caudal peduncles and triggerfish with higher AR caudal fins recruit their caudal fins (U_t low_) at lower speeds than balistoid fishes with wide caudal peduncles and triggerfishes with low AR caudal fins. These relationships are best understood by considering U_t low_ to represent the speed at which caudal fin recruitment is beneficial, rather than the limit of median fin propulsion. Modelling studies have shown that narrow caudal peduncles and high AR caudal fins are more hydrodynamically efficient than wide caudal peduncles and convex caudal fins (Lighthill, 1969, 1970; reviewed in Webb 1982). We likely do not observe a trend between caudal fin AR and U_t low_ within the balistoid fishes as a whole, largely because the filefishes only used the caudal fin for an average of 12% of their total critical swimming performance. This minor contribution of the caudal fin to the overall endurance swimming performance of the filefishes suggests that caudal fin shape is not likely to be evolutionarily specialized for efficient endurance swimming performance in filefishes. Conversely, the triggerfishes, on average, were aided by contribution of the caudal fin during the upper 35% of their total critical swimming performance, suggesting that caudal fin shape may indeed be important for the endurance swimming performance of these triggerfishes at high speeds. This would explain why triggerfishes tend to possess caudal fins of higher ARs than filefishes (2.73 and 2.16 for triggerfishes and filefishes, respectively). The low AR caudal fins and deep caudal peduncles observed in the filefishes in this study are likely more useful for short, quick burst of speed to escape predators than for long, sustained swimming bouts (Weihs, 1973; Webb, 1982). All of these body, median fin and caudal fin traits come together in the *full shape* dataset, and we find a strong correlation between U_t low_ and full shape PC1 and PC2. Specifically fishes with long median fins and relatively wide caudal peduncles (low PC1) and fishes with concave, high AR caudal fins (high PC2) exhibited higher gait transition speeds than fishes with short median fins and narrow caudal peduncles (high PC1) and highly convex caudal fins (low PC2) (Fig. 8B).

Finally, some swimming performance trends are best explained by the percent balistiform locomotion data. Fishes with relatively long and large median fins (regardless of their shape) and fishes with wide caudal peduncles swam using the balistiform locomotion gait alone for a larger percentage of the time. This trend can be explained by the higher maximum power output made possible by large median fins (regardless of fin kinematics) while using the balistiform gait versus increased hydrodynamic efficiency of caudal fin oscillations provided by narrow caudal peduncles. These percent balistiform locomotion trends were some of the cleanest functional morphology trends observed in this study, with few outliers. Each species appears to be fairly specialized for taking advantage of one of these gaits, with balistiform specialists possessing elongate, large median fins, capable of overcoming the large power requirements of swimming at high speeds using the median fins alone, while BCF specialists possess small median fins, incapable of powering high-speed balistiform locomotion, but narrow caudal peduncles capable of facilitating efficient caudal fin oscillations to power high-speed endurance swimming. In order to better understand these functional morphology trends, more work is needed on the energetics and kinematics of each swimming gait across a broad taxonomic balistoid sampling.

### Ecomorphology

The wide range of endurance swimming abilities and gait transition strategies observed between the balistoid species in this study likely has implications for the ecologies of these fishes. All species in this study are reef associated fishes (Randall et al., 1997; Froese and Pauly, 2018), but they don’t all use the reef in the same way. *Odonus niger* and *X. auromarginatus* are largely planktivorous (Fricke, 1980; Randall et al., 1978) and spend large amounts of time swimming well above the reef as they pick plankton from the water column (Fricke, 1980; Meyers, 1991). This behavior explains why these two species have evolved fin and body morphologies suited for providing the highest critical swimming speeds measured in this study. The remaining 11 species can be classified as benthic grazers, as a large portion of their diet is composed of sessile or slow moving benthic organisms (Peristiwady and Geistdoerfer, 1991; Randall, 1955; Randall, 2007; Randall and Hartman, 1968; Hiatt and Strasburg, 1960; Meyer, 1985). Most of these species likely remain close to the shelter of the reef as they nip at algae, small crustaceans, sponges, bivalves, or the coral itself. All filefishes examined in the present study fall into this category, and this lifestyle would likely not require high endurance swimming performance. However, these fishes likely do require the ability to power large bursts of speed to escape from predation into nearby holes in the reef, and this behavior would likely be facilitated by the wide caudal peduncles and large, low aspect ratio caudal fins observed in these filefishes (Weihs, 1973; Webb, 1982). Other benthic grazing species (*B. undulatus, B. conspicillum*, R. *aculeatus, S. bursa* and *S. chrysopterum*) spend much of their time farther from the cover of the reef as they swim along open coral rubble lagoons and feed on more evasive prey such as crabs and even small fishes (Hiatt and Strasburg, 1960; Meyers 1991; Sano et al., 1984;Randall, 1985; Randall, et al. 1997; Vijay Anand and Pillai, 2005). These species likely require some combination of fast, aerobic bursts of speed to catch their prey and escape predators over long distances of open sandy bottoms as well as efficient slow swimming performance to sustain long bouts of searching for benthic prey over the sandy lagoons. These species group in the area of body morphospace defined by narrow caudal peduncles and short median fins and exhibit some of the slowest gait transition speeds (U_t low_) measured in this study (Fig. 8), indicating that they rely heavily on caudal fin contribution to achieve high speed locomotion. Furthermore, research has shown (Korsmeyer et al., 2002) that one species with this body type and these ecological characteristics, *R. aculeatus*, is capable of highly efficient slow swimming using the balistiform gait, as well as sustainable aerobic BCF locomotion at higher speeds. Combined, these trends suggest that the small, short median fins of *R. aculeatus* are sufficient for slow grazing using balistiform locomotion, while the narrow caudal peduncle facilitates efficient high-speed aerobic BCF swimming used to escape predators and chase down elusive prey while swimming over sandy bottom lagoons. In order to determine how well these ecomorphological trends apply to balistoid fishes as a whole, more research is required on the morphometrics and ecologies of a larger, phylogenetically informed sampling of the superfamily Balistoidea.

## Acknowledgments

We thank Mr. Hiroaki Hayashi and Dr. Hiroshi Senou of the Kanagawa Prefectural Museum of Natural History (KPM) for measuring and providing high resolution photos of all KPM specimens used in this study. We thank Douglas Nelson of UMMZ and Dr. Caleb McMahan and Susan Mochel of the FMNH for providing lab space and specimens. We thank Isaac Krone, Drs. Aaron Olsen, Charlene McCord and Andrew Hipp for advice concerning geometric morphometric analyses. We thank Dr. Brad Wright for developing the swimming protocol used in this study. Finally, we thank the following museums for access to specimens or photographs used in morphometric analyses: KPM, FMNH, UMMZ, BPBM, NMNH and ROM.

## Literature Cited

Adams, D. C., Collyer, M. L., Kaliontzopoulou, A. and Sherratt, E. (2017). Geomorph: Software for geometric morphometric analyses. R package version 3.0.5. https://cran.rproject.org/package=geomorph.

Alexander, R. (1989). Optimization and gaits in the locomotion of vertebrates. Physiol. Rev. 69, 1199–1227.

Alsop, D. H. and Wood, C. W. (1997). The interactive effects of feeding and exercise on oxygen consumption, swimming performance and protein usage in juvenile rainbow trout (*Oncorhynchus mykiss*). J. Exp. Biol. 200, 2337–2346.

Benjamini, Y. and Hochberg, Y. (1995). Controlling the false discovery rate: a practical and powerful approach to multiple testing. J. R. Stat. Soc. B. 57, 289–300.

Blake, R. W. (1978). On balistiform locomotion. J. Mar. Biol. Assoc. UK. 58, 73–80.

Breder, C. M. (1926). The locomotion of fishes. Zoologica-New York. 4, 159–256.

Brett, J. (1964). The Respiratory Metabolism and Swimming Performance of Young Sockeye Salmon. J. Fish Res. Board Can. 21, 1183–1226.

Bushnell, D. M. and Moore, K. J. (1991). Drag reduction in nature. Ann. Rev. Fluid. Mech. 23, 65–79.

Cannas, M. Schaefer, J., Domenici, P. and Steffensen, J. (2006). Gait transition and oxygen consumption in swimming striped surfperch *Emiotoca lateralis* Agassiz. J. Fish Biol. 66, 1612–1625.

Dornburg, A., Santini, F. and Alfaro, M. E. (2008). The influence of model averaging on clade posteriors: an example using the triggerfishes (Family Balistidae). Syst. Biol. 57, 905–919.

Dornburg, A., Sidlauskas, B., Santini, F., Sorenson, L., Near, T. J. and Alfaro, M. E. (2011). The influence of an innovative locomotor strategy on the phenotypic diversification of triggerfish (Family Balistidae). Evolution 65, 1912–1926.

Drucker, E. G. and Jensen, J. S. (1996). Pectoral fin locomotion in the striped surfperch. II. Scaling swimming kinematics and performance at a gait transition. J. Exp. Biol. 199, 2243–2252.

Farlinger, S. and Beamish, F. W. H. (1977). Effects of time and velocity increments on the critical swimming speed of largemouth bass (*Micropterus salmoides*). Trans. Am. Fish. Soc. 106, 436–439.

Feilich, K. L. (2016). Correlated evolution of body and fin morphology in the cichlid fishes. Evolution. 70, 2247–2267.

Feilich, K. L. (2017). Swimming with multiple propulsors: measurement and comparison of swimming gaits in three species of neotropical cichlids. J. Exp. Biol. 220, 4242–4251.

Fricke, H. W. (1980). Mating systems, maternal and biparental care in triggerfish (*Balistidae*). Z. Tierphychol. 53, 105–122.

Froese, R. and Pauly, D. (2018). FishBase. <http://www.fishbase.org>.

Fulton, C. J. and Bellwood, D. R. (2004). Wave exposure, swimming performance, and the structure of tropical and temperate reef fish assemblages. Mar. Biol. 144, 429–437.

Hale, M. E., Day, R. D., Thorsen, D. H. and Westneat, M. W. (2006). Pectoral fin coordination and gait transitions in steadily swimming juvenile reef fishes. J. Exp. Biol. 209, 3708–3718.

Hiatt, R. W. and Strasburg, D. W. (1960). Ecological relationships of the fish fauna on coral reefs of the Marshall Islands. Ecol. Monogr. 30, 65–127.

Hu, T., Wang, G., Shen, L. and Li, F. (2006). A novel conceptual fish-like robot inspired by *Rhinecanthus aculeatus*. 2006 9th International Conference on Control, Automation, Robotics and Vision, Singapore. pp. 1–5.

Hutchins, J. B. and Swainston, R. (1985). Revision of the Monacanthid fish genus *Brachaluteres*. Rec. West. Aust. Mus. 1, 57–78.

Karpouzian, G., Spedding, G. and Cheng, H. K. (1990). Lunate-tail swimming propulsion. Part 2. Performance analysis. J. Fluid. Mech. 210, 329–351.

Korsmeyer, K. E., Steffensen, J. F. and Herskin, J. (2002). Energetics of median and paired fin swimming, body and caudal fin swimming, and gait transition in parrotfish (*Scarus schlegeli*) and triggerfish (*Rhinecanthus aculeatus*). J. Exp. Biol. 205, 1253–1263.

Kolok, A. S. (1999). Interindividual variation in the prolonged locomotor performance of ectothermic vertebrates: a comparison of fish and herpetofaunal methodologies and a brief review of the recent fish literature. Can. J. Fish. Aquat. Sci. 56, 700–710.

Lighthill, M. J. (1969). Hydromechanics of aquatic animal propulsion. Annu. Rev. Fluid Mech. 1, 413–446.

Lighthill, M. J. (1970). Aquatic animal propulsion of hydrodynamical efficiency. J. Fluid Mech. 44, 265–301.

Lighthill, M. J. and Blake, R. (1990). Biofluiddynamics of balistiform and gymnotiform locomotion. Part 1. Biological background, and analysis by elongated-body theory. J. Fluid Mech. 22, 183–207.

Loofbourrow, H. (2009). Hydrodynamics of balistiform swimming in the Picasso triggerfish, Rhinecanthus aculeatus. Master’s thesis, University of British Columbia, Vancouver, Canada.

MacLeod, N. (2009). Form & shape models. Palaeontology Newsletter. 18, 1–11.

McCord, C. L. and Westneat, M. W. (2016). Phylogenetic relationships and the evolution of BMP4 in triggerfishes and filefishes (Balistoidea). Mol. Phylogenet. Evol. 94, 397–409.

Millard, S. P. (2013). EnvStats: An R Package for Environments Statistics. New York: Springer.

Meyer, D. L. (1985). Evolutionary implications of predation on recent comatulid crinoids from the Great Barrier Reef. Paleobiology. 11, 154–164.

Meyers, R. F. (1991). Micronesian Reef Fishes: A Practical Guide to the Identification of the Coral Reef Fishes of the Tropical Central and Western Pacific. 2nd ed. Gaum, USA: Coral Graphics.

Nursall, J. T. (1958). The caudal fin as a hydrofoil. Evolution. 12, 116–120.

Olsen, A. M. (2017). Feeding ecology is the primary driver of beak shape diversification in waterfowl. Funct. Ecol. 31, 1985–1995.

Olsen A. M. and Westneat, M. W. (2015). StereoMorph: an R package for the collection of 3D landmarks and curves using a stereo camera setup. Methods Ecol. Evol. 6, 351–356.

Orme, C. D. L., Freckleton, R. P., Thomas, G. H., Petzoldt, T., Fritz, S. A. and Isaac, N. (2013). caper: comparative analyses of phylogenetics and evolution in R. R package version0.5.2. https://CRAN.R-project.org/package=caper.

Peristiwady, T. and Geistdoerfer, P. (1991). Biological aspects of *Monacanthus tomentosus* (Monacanthidae) in the seagrass beds of Kotania Bay, West Seram, Moluccas, Indonesia. Mar. Biol. 109, 135–139.

R Core Team. (2016). R: a language and environment for statistical computing. Vienna, Austria: R Foundation for Statistical Computing.

Randall, J. E. (1955). Fishes of the Gilbert Islands. Atoll Research Bulletin. 47, 1–243.

Randall, J. E. (1964). A revision of the filefish genera *Amanses* and *Cantherhines*. Copeia. 1964, 331–361.

Randall, J. E. (1985). Guide to Hawaiian Reef Fishes. Newtown Square, PA: Harrowood Books.

Randall, J. E. (2007). Reef and Shore Fishes of the Hawaiian Islands. Honolulu, HI: University of Hawai’i Sea Grant College Program.

Randall, J. E. and Hartman, W. (1968). Sponge-feeding fishes of the West Indies. Mar. Biol. 1, 216–225.

Randall, J. E., Matsuura, K. and Zama, A. (1978). A revision of the triggerfish genus *Xanthinichthys*, with description of a new species. B. Mar. Sci. 28, 688–706.

Randall, J. E., Allen, G. R. and Steene, R. C. (1997). Fishes of the Great Barrier Reef and Coral Sea. Revised and Expanded Edition. Honolulu, HI: University of Hawaii Press.

Revell, L. J. (2012) phytools: An R package for phylogenetic comparative biology (and other things). Methods Ecol. Evol. 3, 217–223.

Rouleau, S., Glémet, H. and Magnan, P. (2010). Effects of morphology on swimming performance in wild and laboratory crosses of brook trout ecotypes. Funct. Ecol. 24, 310–321.

Sano, M., Shimizu, M. and Nose, Y. (1984). Food habits of Teleostean reef fishes in Okinawa Island, Southern Japan. Tokyo, Japan: University of Tokyo Press.

Santini, F., Sorenson, L. and Alfaro, M. E. (2013). A new multi-locus timescale reveals the evolutionary basis of diversity patterns in triggerfishes and filefishes (Balistidae, Monacanthidae; Tetraodontiformes). Mol. Phylogenet. Evol. 69, 165–176.

Sfakiotakis, M., Lane, D. M. and Davies, J. B. C. (1999). Review of fish swimming modes for aquatic locomotion. IEEE J. Ocean. Eng. 24, 237–252.

Sprinkle, B., Bale, R., Bhalla, A. P. S., MacIver, M. A. and Patankar, N. A. (2017). Hydrodynamic optimality of balistiform and gymnotiform locomotion. European Journal of Computational Mechanics. 26, 31–43.

Svendsen, J. C., Tudorache, C., Jordan, A. D., Steffensen, J. F., Aarestrup, K. and Domenici, P. (2010). Partition of aerobic and anaerobic swimming costs related to gait transitions in a labriform swimmer. J. Exp. Biol. 213, 2177–2183.

Vijay Anand, P. E. and Pillai, N. G. K, (2005). Community organization of coral reef fishes in the rubble sub-habitat of Kavaratti Atoll, Lakshadweep, India. J. Mar. Biol. Assoc. India 47, 7782.

Vogel, S. (1994). Life in Moving Fluids: The Physical Biology of Flow. 2nd ed. Princeton, NJ: Princeton University Press. pp. 224–233.

Wainwright, P. C., Bellwood, D. R. and Westneat, M. W. (2002). Ecomorphology of locomotion in labrid fishes. Environ. Biol. Fish. 65, 47–62.

Walker, J. A. and Westneat, M. W. (2002). Performance limits of labriform propulsion and correlates with fin shape and motion. J. Exp. Biol. 205, 177–187.

Webb, P. W. (1982). Locomotor patterns in the evolution of Actinopterygian fishes. Am. Zool. 22, 329–342.

Webb, P. W. (1984). Body form, locomotion and foraging in aquatic animals. Am. Zool. 24, 107–120.

Weihs, D. (1973). The mechanism of rapid starting of slender fish. Biorheology. 10, 343–350.

Whoriskey, F. G. and Wooton, R. J. (1987). The swimming endurance of threespine sticklebacks, *Gasterosteus aculeatus* L., from the Afon Rheidol, Wales. J. Fish Biol. 30, 335–339.

Winterbottom, R., Emery, A., and Holm, E. (1989). An Annotated Checklist of the Fishes of the Chagos Archipelago, Central Indian Ocean. Toronto, Canada: Royal Ontario Museum.

Wright, B. (2000). Form and function in aquatic flapping and propulsion: morphology, kinematics, hydrodynamics, and performance of the triggerfishes (Tetraodontiformes: Balistidae). PhD thesis, University of Chicago, Chicago, IL.

Xin, Z. Q. and Wu, C. J. (2013). Shape optimization of the caudal fin of the three-dimensional self propelled swimming fish. Sci. China Phys. Mech. 56, 328–339.

